# Development and Characterization of a low intensity vibrational system for microgravity studies

**DOI:** 10.1101/2023.11.20.567870

**Authors:** Omor M. Khan, Will Gasperini, Chess Necessary, Zach Jacobs, Sam Perry, Jason Rexroat, Kendall Nelson, Paul Gamble, Twyman Clements, Maximilien DeLeon, Sean Howard, Anamaria Zavala, Mary Farach-Carson, Elizabeth Blaber, Danielle Wu, Aykut Satici, Gunes Uzer

**Author notes:** **Corresponding authors:** Aykut Satici PhD, Gunes Uzer PhD Boise State University Department of Mechanical & Biomedical Engineering 1910 University Drive, MS-2085 Boise, ID 83725-2085. Funding support: AG059923, P20GM109095, P20GM103408, NSF1929188 and, NSF 2025505.

## Abstract

The advent of extended-duration human spaceflight demands a better comprehension of the physiological impacts of microgravity. One primary concern is the adverse impact on the musculoskeletal system, including muscle atrophy and bone density reduction. Ground-based microgravity simulations have provided insights, with vibrational bioreactors emerging as potential mitigators of these negative effects. Despite the potential they have, the adaptation of vibrational bioreactors for space remains unfulfilled, resulting in a significant gap in microgravity research. This paper introduces the first automated low-intensity vibrational (LIV) bioreactor designed specifically for the International Space Station (ISS) environment. Our research covers the bioreactor’s design and characterization, the selection of an optimal linear guide for consistent 1-axis acceleration, a thorough analysis of its thermal and diffusion dynamics, and the pioneering use of BioMed Clear resin for enhanced scaffold design. This advancement sets the stage for more authentic space-based biological studies, vital for ensuring the safety of future space explorations.

## Introduction

The 21st century has witnessed notable advancements in the field of space exploration, as both private and public sectors have made substantial efforts to expand the boundaries of human presence beyond Earth^1–9^. In order to ensure the continued viability of long-term spaceflight, it is critical to address the physiological challenges that arise from microgravity as mission duration and reach increase^2,4,9,10^. Microgravity poses a unique challenge to astronaut health, particularly affecting the musculoskeletal system. Muscle atrophy and bone density reduction are among the most pressing health issues faced during extended missions, necessitating innovative solutions to mitigate these effects^10–19^.

Ground-based studies have provided valuable insights into the challenges of microgravity. Simulated microgravity experiments, particularly those employing low- and high-intensity vibrational bioreactors, have shown promise in mitigating some of the adverse physiological effects of simulated microgravity^16,17,19–27^. In addition, pharmacological treatments for bone loss resulting from exposure to microgravity are also being tackled and researched in both real and simulated microgravity^10,11,28^.

Nevertheless, the full mechanism and impact of microgravity on the human body, especially in real space conditions, are still not entirely understood^12^. Astronauts aboard the International Space Station (ISS) undergo rigorous exercise regimens to combat the adverse effects of living in space^10–12,17–19,28^. While these exercises help, they don’t completely negate the physiological changes induced by microgravity^12,19^.

These challenges highlight the crucial necessity of conducting direct cellular studies in a microgravity setting and evaluating the possible advantages of interventions such as vibrational bioreactors in the context of space exploration. While vibrational bioreactors have shown promise on Earth^14–17,21–26^, there is a conspicuous absence of such technology tailored for use in space^19^, especially one that meets the stringent standards of the ISS.

Our research is motivated by the current limitations in understanding the effects of microgravity at cellular level. The primary aim of our study is to design and characterize an automated low-intensity vibrational (LIV) bioreactor for studying microgravity. This bioreactor is tailored to meet the stringent requirements of the (ISS). Until now, there have been no vibrational bioreactors in space. With the introduction of our LIV bioreactor, we aim to bridge this gap, enabling a direct comparison between experiments conducted in space and those on Earth. This endeavor is crucial for advancing our understanding and ensuring the safety and sustainability of future human space exploration.

### Experimental Design

The objective of our research was to engineer a low-intensity vibrational (LIV) bioreactor suitable for microgravity conditions aboard the (ISS). Our design had to conform to the CubeLab Interface Control Document (ICD)^29^ standards, ensuring compatibility with the ISS’s operational environment. We utilized the CubeLab 9U module (**Supplementary Figure 2**), provided by Space Tango^30^, which offers a standardized platform for microgravity experiments^2,31^, allowing for a secure and controlled setting for our bioreactor.

In a precursor study within our laboratory, we developed a 3D bone analog scaffold utilizing gyroid shapes to mimic the complex architecture of bone tissue^32^. For the current bioreactor design, these scaffolds are integral to our assessment, serving as a platform to implement and evaluate the bioreactor’s capacity to support biological studies. The focus of our investigation is the application of vibrational stimuli to bone cells, addressing the challenge of bone density reduction—a significant concern for astronauts in microgravity conditions. This approach aims to leverage the mechanical signals provided by our bioreactor to enhance bone cell function and mitigate the effects of space-induced osteopenia.

For the delivery of precise vibrational stimuli, we selected the piezoelectric actuator (P-841.30, Physik Instrumente (PI), Karlsruhe, Germany) and its closed-loop controller (E-610.S0, Physik Instrumente (PI), Karlsruhe, Germany), chosen for their compliance with the CubeLab’s size, weight, and power constraints. These components were tasked with providing a consistent vibration regime at 90Hz with a 0.7g peak-to-peak (1g=9.81 m/s^2^) acceleration, ensuring the application of low-intensity vibrations to the cells.

Our methodology began with the selection of an appropriate linear guide to facilitate precise one-axis acceleration. This selection process was informed by evaluating multiple linear guides against our design criteria.

Next, we conducted thermal and vibrational analyses of the bioreactor’s behavior using both COMSOL simulations and empirical testing to ensure that the applied vibrations were within the operational parameters and did not produce too much heat, compromising cell viability. Additionally, we made sure that any resonant frequencies of the system are outside of its operational range so that they do not interfere with the bioreactor’s function.

To validate the bioreactor’s compatibility with cellular studies, we performed XTT and Live/Dead assays to confirm that the bioreactor design did not adversely affect cell viability. These assays were critical in demonstrating the biocompatibility of our system.

Finally, we addressed the challenge of media exchange within the CubeWells. A diffusion study^33^, supported by COMSOL simulations and visual assessments using a real-time camera setup, was conducted to ascertain the efficiency of media introduction and distribution within the wells, ensuring that all cells received adequate nutrients without the risk of shear-stress-induced damage.

### Methods and Data Collection

In order to drive the our piezoelectric actuator’s controller (E-610.S0) we needed to generate a 90Hz 0-10V signal from a Teensy microcontroller (PJRC, TEENSY41), which only provides a 0-3.3V PWM output. A custom amplifier circuit was designed with a Sallen-Key^34^ second-order lowpass filter topology (**Supplementary Figure 1**). The filter was tuned to isolate a 90 Hz sine wave from a modulated PWM signal, removing the extraneous frequencies. This process produced a clean sine wave to drive the piezoelectric actuator, providing the precise vibrational stimulus required for the biological experiments.

### Linear guide selection

To select a linear guide capable of delivering precise one-axis acceleration, we conducted performance evaluations on three different models. Using LabVIEW software, we measured the acceleration profiles of each guide with a MEMSIC CXL04GP3 accelerometer (**Supplementary Figure 3**). The guides were tested for their ability to sustain a 0.7g peak-to-peak acceleration at a frequency of 90 Hz over a 20-minute duration. The objective was to identify a guide that would restrict vibrations to the intended axis, ensuring accurate delivery of mechanical stimuli to the samples. The tested guides (**Figure 1**) included: a custom-built roller-based guide (SS2EB, Misumi, Schaumburg, IL), known for smooth motion; a low-maintenance plastic linear guide (N Prism Preloaded, Igus Inc., Rumford, RI); and a Schneeberger roller-based linear guide (NKL 2-95, Schneeberger Inc., Woburn, MA).

**Figure 1:**
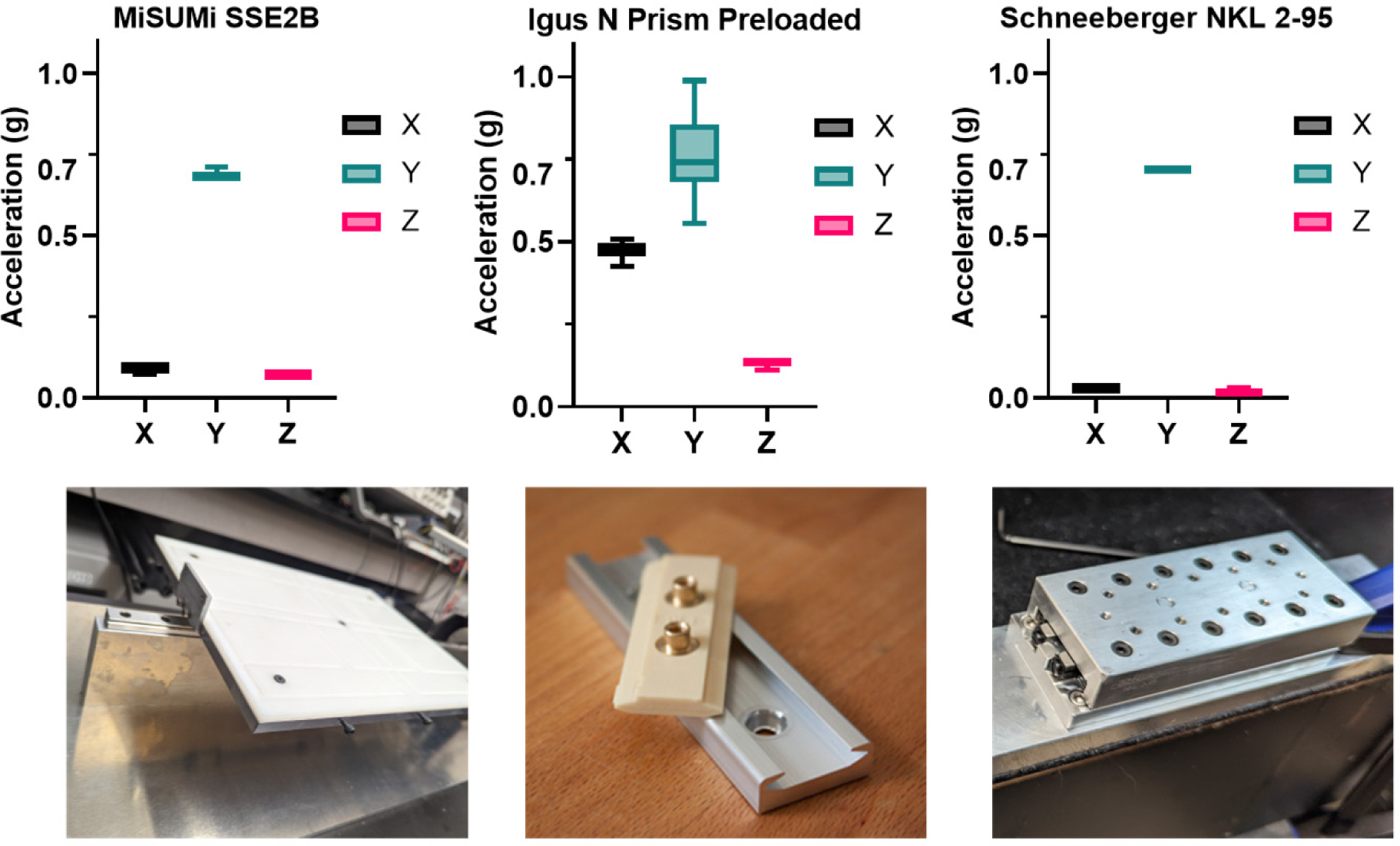
Comparison of peak-to-peak accelerations for three different linear guides under 90Hz, 0.7g peak-to-peak vibration. The guides tested were an N-Prism linear guide, a Schneeberger linear guide, and a Misumi linear guide. Box and violin plots represent the distribution of peak-to-peak accelerations in the X, Y, and Z axes over a 20-minute test period. The Schneeberger guide demonstrated the most consistent performance, with a peak-to-peak acceleration of 0.7g in the Y-axis and minimal vibration translation to the X and Z axes.

### Thermal Characterization

The thermal performance of the LIV bioreactor was evaluated to determine its suitability for sustaining biological experiments within the CubeLab module. The fully assembled bioreactor, including the piezoelectric actuator, closed-loop controller, and the selected linear guide, was installed inside the CubeLab. Thermocouples (Type K) were positioned: directly inside the controller housing, above the actuator, adjacent to the biological sample area, and within the external ambient environment of the CubeLab (**Figure 2**).

**Figure 2:**
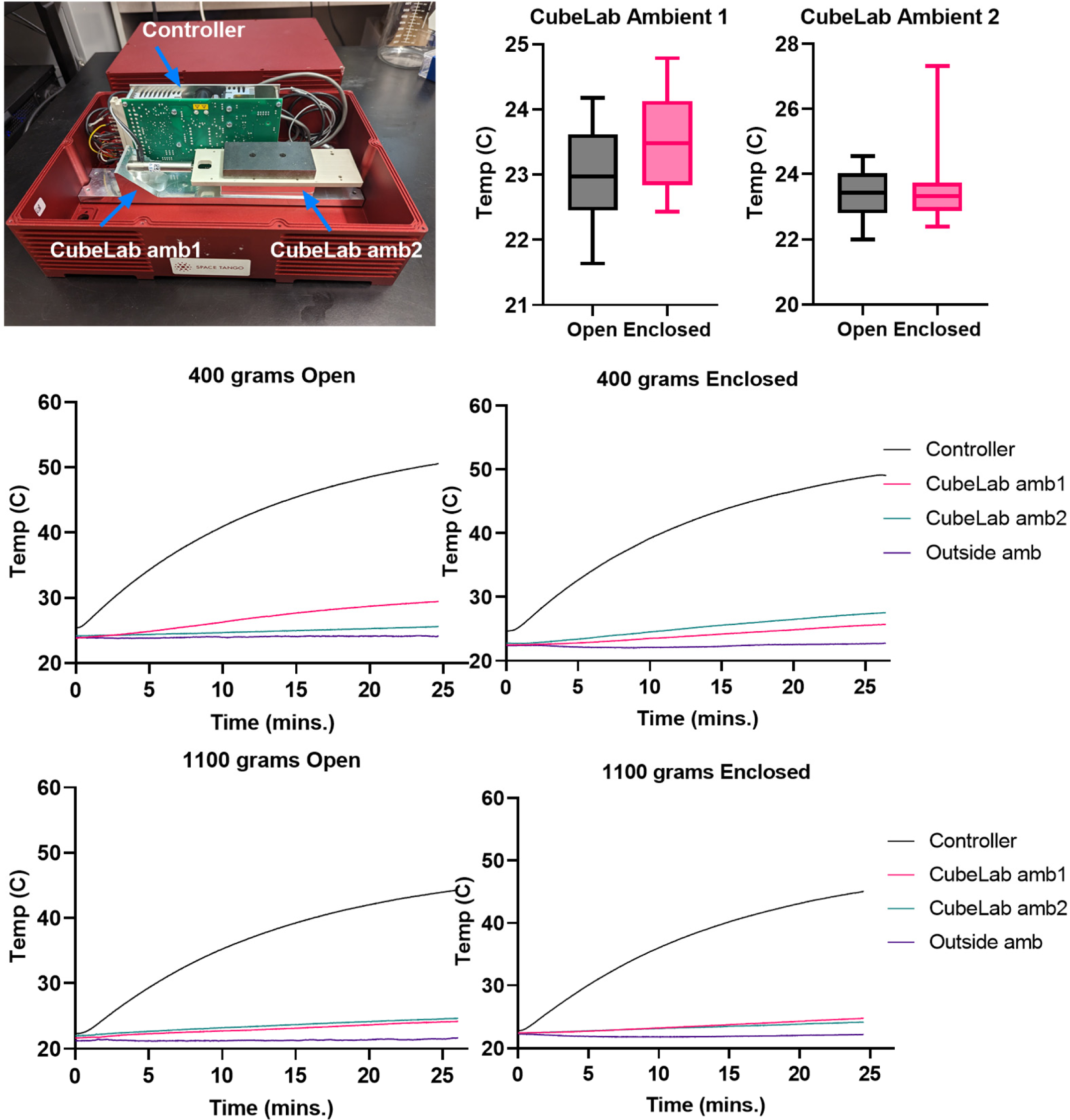
Differential Heating Effects on the Vibrational Bioreactor and Surrounding Environment. This figure illustrates the varying temperature profiles observed during a 25-minute vibrational experiment conducted within a closed and open-lid system. Internal and external thermocouples recorded the temperature of the vibrational controller (rising to 59°C) and the ambient air within and outside the enclosure (increasing from 21°C to 25°C). The results confirm that despite the significant heat generated by the vibrational actuator, the ambient conditions remained within a safe range for cell viability.

LabVIEW software with the NI 9211 DAQ card (National Instruments, Austin, TX, USA) was utilized for both control of the vibrational input and data acquisition of the thermal output. The system was tested with two different payload conditions, 400 grams and 1100 grams, to simulate varying experimental loads. The operational test consisted of a 25-minute vibration cycle at 0.7g peak-to-peak acceleration and a frequency of 90Hz.

Thermal characterization was conducted under two distinct CubeLab configurations: an ‘Open’ setup with the top cover removed and a ‘Closed’ setup where the module was entirely sealed. These tests were performed without the integration of active cooling systems to assess the bioreactor’s inherent thermal management capabilities under operational conditions. Data collected during these tests were analyzed to understand the thermal dynamics of the system, with particular attention to the temperature stability and heat dissipation efficiency in both ‘Open’ and ‘Closed’ states.

### Vibration Characterization

The vibrational characteristics of our bioreactor were analyzed through both simulation and experimental testing. In the simulation phase, COMSOL was used to study the system’s dynamics. A detailed CAD model of the bioreactor, which included the metal base, actuator, linear guide, and payload holder, was integrated into the simulation environment. We follows a simple, conservative approach in our analysis by assuming that, the material for the piezoelectric actuator was the same as its housing - standard stainless steel. This assumption allowed us to focus on the primary vibration modes crucial for the bioreactor’s functionality in microgravity, without the complexities of modeling the piezoelectric properties. This setup, which mirrored real-world constraints and was executed with a standard mesh configuration, provided a robust and efficient method to assess the vibrational dynamics of the bioreactor.

Concurrently, experimental tests were conducted to validate the simulation results and further understand the bioreactor’s vibrational behavior. The system was subjected to vibrations at a targeted 0.4g peak-to-peak acceleration across a frequency range of 50 Hz to 500 Hz, with finer increments around the critical 90 Hz range. Subsequent tests aimed for a 0.7g peak-to-peak vibration; however, due to the actuator’s stroke constraints, only at frequencies from 80 Hz to 150 Hz.

### Scaffold Viability

Cell viability on SLA-printed scaffolds made from BioMed Clear resin was assessed using XTT assays for quantitative analysis and Live/Dead imaging for qualitative evaluation. The resin was selected for its precision in creating complex structures. Scaffolds were integrated with a hydrogel to improve the biomimetic quality for cell culture studies.

#### Cell Culture and Seeding

Murine mesenchymal stem cells (MSCs) were employed for this study. These cells were encapsulated within a hydrogel^35(p3)^ composed of Thiolated Hyaluronic Acid (HA) (Advanced Biomatrix, #GS22F), functionalized with specific amino sequences: PQ (GGGPQ↓IWGQGK concentrated at 44.973 mgs/mL) and RGD (GRGDS concentrated at 73.7mgs/mL) with a volume ratio of 4 HA:1 RGD: 1PQ. The encapsulated cells were then seeded into gyroid-shaped scaffolds, designed to mimic the bone marrow microenvironment. These scaffolds were fabricated using the BioMed Clear resin from Formlabs and were SLA printed using a Form 2 printer.

#### Osteogenic Media Preparation

The osteogenic media used for cell culture was prepared using α-MEM, supplemented with 10% FBS, 100 U/mL penicillin, 100 µg/mL streptomycin, Vitamin C at 50ug/ml, and beta-glycerophosphate ranging from 10mM-20mM.

#### Vibration Regime

Post-seeding, the scaffolds were transferred to a 96-well plate containing the osteogenic media. The entire setup was then placed in an incubator. A distinct vibration regime was implemented, wherein the samples were subjected to a consistent vibration of 0.7g peak-to-peak at 90 Hz for 20 minutes. This vibration was administered twice daily, with a 2-hour interval between each session.

#### XTT Assay

On the first and fourth day, an XTT assay was performed to assess cell viability. The assay utilized two primary reagents: an electron mediator solution and the XTT developer reagent (Cayman Chemical, #10010200). Following the manufacturer’s protocol, the samples were incubated for approximately 2 hours at 37°C. The absorbance of the assay wells was then measured using a microtiter plate reader, capturing values between 400-500 nm wavelengths.

#### Live/Dead Imaging

On the fifth day, live/dead imaging was conducted. The cells were stained using Calcein AM (green) to identify live cells, Propidium Iodide (red) to mark dead cells, and Hoechst 33342 (blue) for nuclear staining according to the manufacturer’s specifications (EMD Millipore #CBA415). The stained samples were then visualized under a fluorescence microscope. While no quantitative data was extracted from these images, they provided a visual representation of the ratio between live and dead cells, offering insights into cell health and viability.

#### Controls and Replicates

The study employed both vibrated and non-vibrated samples as experimental and control groups (+LIV/-LIV), respectively. Each week’s samples (n=4/week for 2 weeks) were biological replicas, with fresh samples prepared for the subsequent week following the identical protocol as the initial week.

### Diffusion dynamics within the CubeWells

To ensure consistent cell culture conditions within the custom-designed CubeWells by Space Tango, a diffusion study was conducted. Initial tests verified the CubeWells’ structural integrity through a visual leak assessment. The primary objectives of the diffusion study^33^ were to evaluate the efficiency of a miniaturized pump in introducing new media into the CubeWell and to confirm uniform media distribution across the well. This was achieved through both COMSOL simulations and real-time camera observations.

#### COMSOL Simulation

To predict the diffusion dynamics within the custom-designed CubeWell system, a multiphysics simulation was conducted using COMSOL. The study combined two primary modules: Laminar Flow and Transport of Diluted Species. The Laminar Flow module was employed to simulate the fluid flow into the CubeWell, while the Transport of Diluted Species module was used to visualize the influx of new media, set at a concentration of 1 mol/m^3^. The simulation was constrained by defining the inlet at the top and outlet at the bottom of the CubeWell (**Figure 5b**), along with the boundary walls, ensuring a realistic representation of the media exchange process.

**Figure 3:**
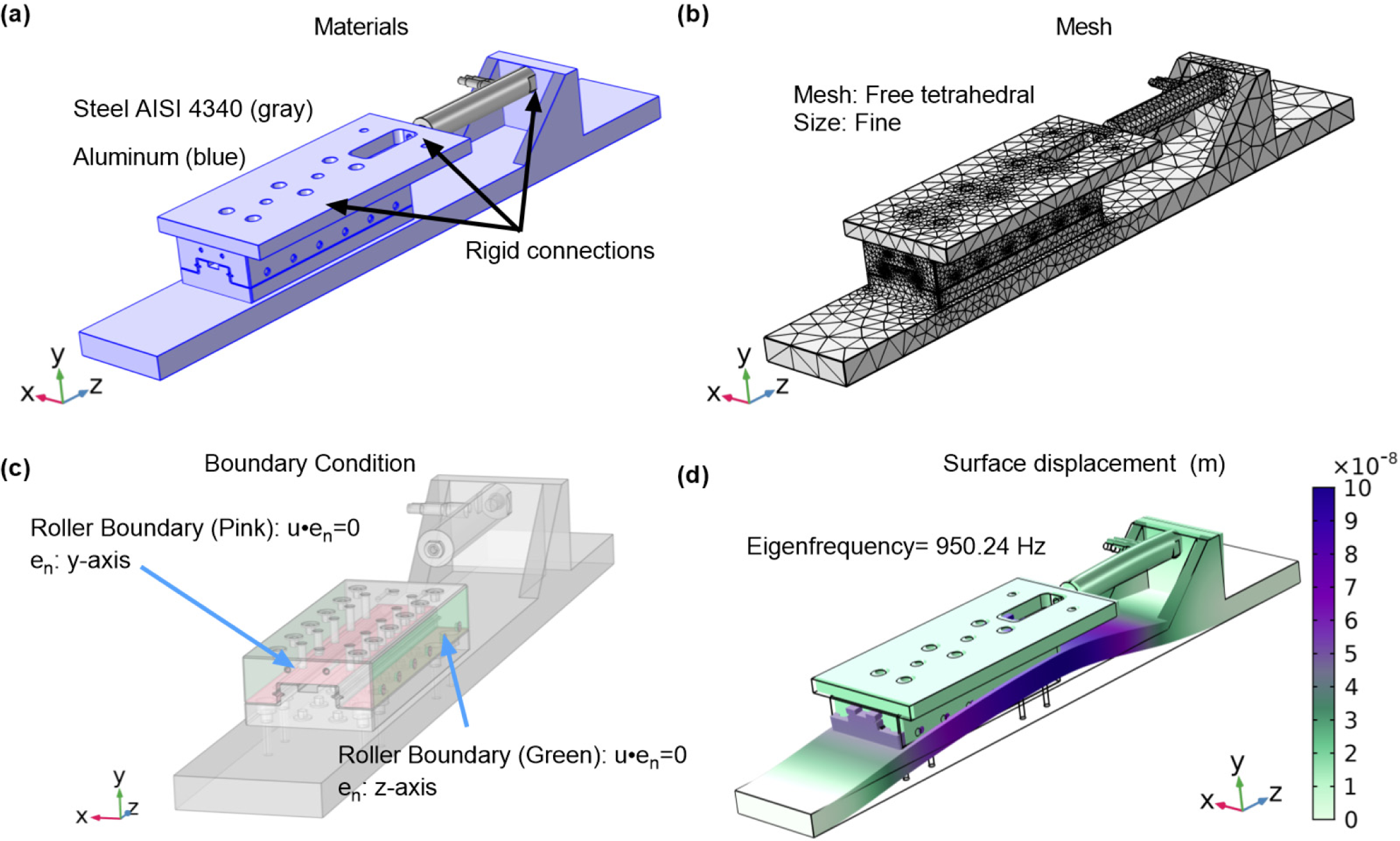
Vibrational analysis of the bioreactor using COMSOL Multiphysics. **(a)** The bioreactor assembly with components materials assigned. **(b)** The applied mesh ensuring resolution for accurate mode shape determination. **(c)** Roller boundary conditions are applied to simulate linear guide constraints **(d)** The first mode shape at 950.24 Hz, demonstrating primary deformation.

**Figure 4:**
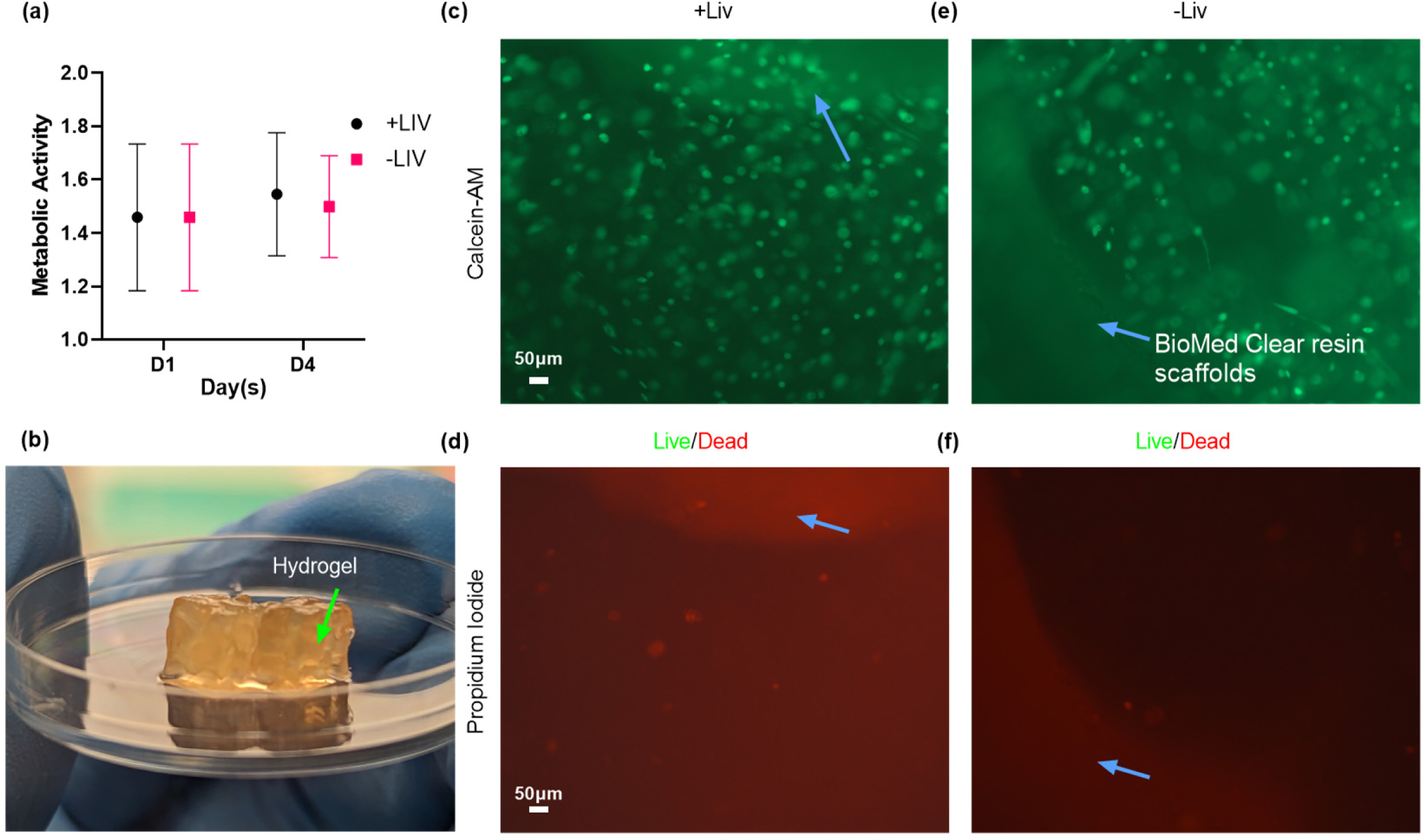
Analysis of Stem Cell Viability and Metabolic Activity in SLA-Printed Scaffolds. This figure presents the results of a 4-day experiment investigating the survival and metabolic activity of stem cells grown in SLA-printed scaffolds (blue arrows). **(a)** Metabolic activity was measured using an XTT assay at day 1 (mean = 1.4, SD = 0.23) and at day 4 (mean = 1.5, SD = 0.19), showing a slight increase over time despite high standard deviations due to a small sample size (n = 4 per group). Analysis showed no significance between groups and days. Concurrently, **(c-f)** Live/Dead staining and imaging highlighted a predominance of viable (green) cells over dead ones, reinforcing the potential for successful cell growth within these scaffolds.

**Figure 5:**
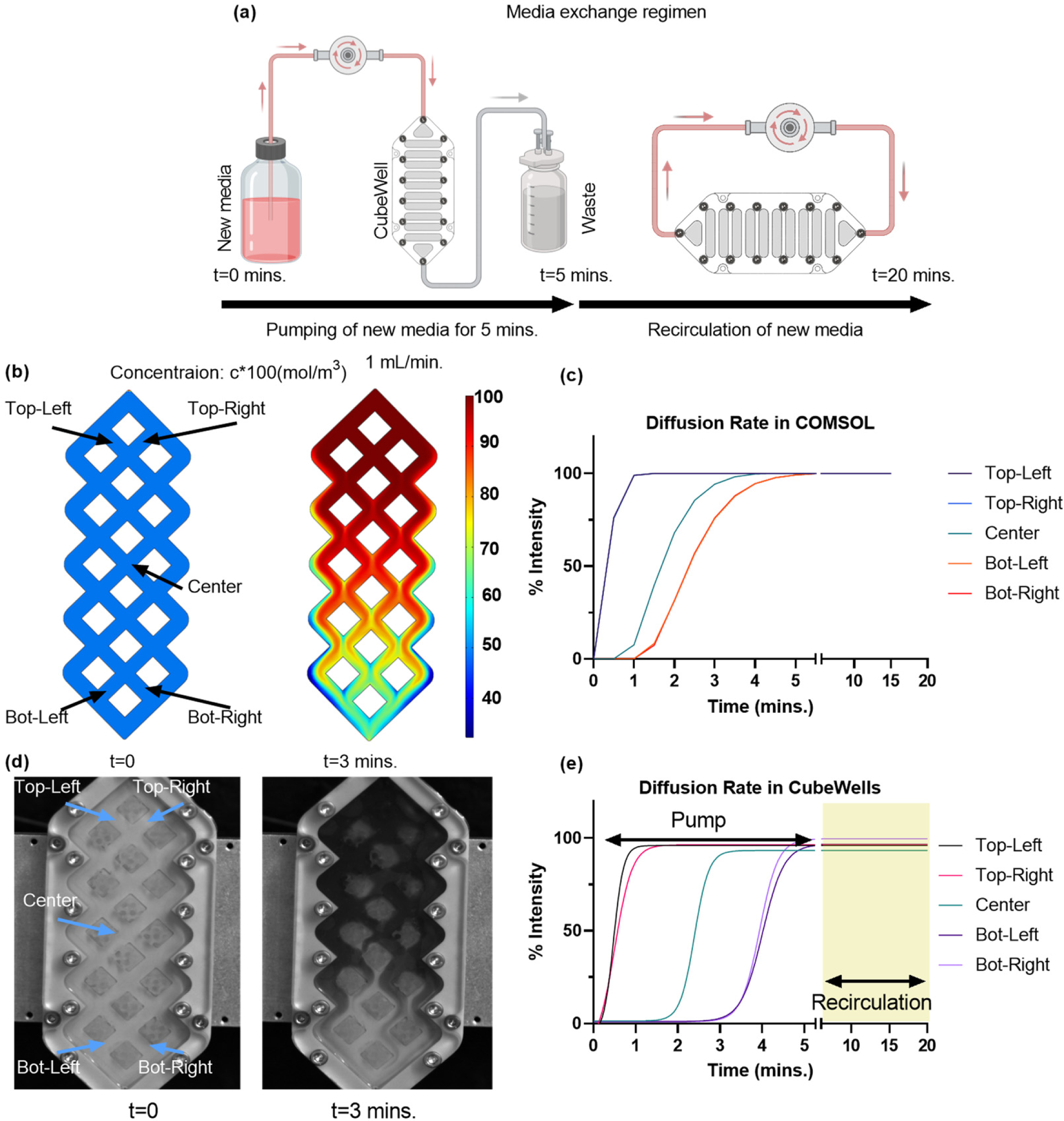
Diffusion study of custom well plate for cell culture. **(a)** For most efficient volume change, a 5-minute pump followed by a 15 min. recirculation is implemented in the media change regimen for the real experiment **(b)** COMSOL simulation depicting media concentration profile and its **(c)** quantitative analysis. **(d)** Real experiment replicating the COMSOL simulation using dye as new media for visualization and **(e)** intensity values of the real experiment showcasing the concentration gradient within the 5 min. pump and 15 min. recirculation in highlighted region

#### Experimental Setup

For the in-vitro experiment, the CubeWells were pre-filled with 8 mL of water. Within this environment, SLA-printed scaffolds made from BioMed Clear resin were introduced. These scaffolds, measuring 5x5x5mm, were encapsulated in hydrogels (**Figure 4b**), mimicking the cellular constructs intended for subsequent cell culture experiments.

#### Media Exchange and Observation

To simulate the introduction of fresh media, a mixture of water and Royal Blue Icing color dye (Wilton, #17111150) was prepared. This simulated medium was then pumped into the CubeWell at a consistent rate of 1 mL/min. The entire process of media exchange, including the diffusion of the dye and fluid movement within the CubeWells, was captured using a high-speed camera (Photron FASTCAM MINI UX50) set at 1000 frames per second (fps). This recording spanned a duration of 5 minutes, providing a detailed visual account of the diffusion dynamics.

#### Data Analysis

Post-recording, the captured footage was analyzed to quantify diffusion within the CubeWell system. Five distinct locations within the CubeWell were selected for this analysis. At these points, the intensity values of the dye were measured over time, offering a quantitative perspective on the diffusion rates and patterns.

## Results

### Linear guide selection

The vibration experiment results (**Figure 1)** confirmed the Schneeberger NKL 2-95 linear guide’s superior performance, with a 95.47% and 52.60% more consistent y-axis acceleration than the N preloaded prism carriage and the custom-built laboratory guide, respectively. The Schneeberger guide maintained a stable acceleration profile with lower mean values (X: 0.031g, Y: 0.704g, Z: 0.018g) and standard deviations (X: ±0.000g, Y: ±0.005g, Z: ±0.006g), indicating its precise control over vibrational input. In comparison, the N preloaded prism carriage showed higher variability, particularly in the x-axis (0.473g±0.024) and z-axis (0.133g±0.008), while the laboratory guide using Misumi roller guides presented intermediate stability (X: 0.089g±0.007, Y: 0.684g±0.011, Z: 0.071g±0.000). All guides met the experimental requirement of 0.7g peak-to-peak acceleration in the y-axis, demonstrating their adequacy for the intended vibrational studies.

### Thermal profile in the CubeLab environment

Thermal profile (**Figure 2**) of each component measured in the CubeLab maintained a steady temperature in different conditions (Open and Closed), with variations within ±0.80°C over the 25-minute LIV period. In the ‘Open’ CubeLab configuration, where the top cover was removed, the temperature readings indicated a gradual increase over time in the controller and the ambient air temperature 1 and 2 in the CubeLab. Although the ambient air temperature increased, the temperatures for ambient 1 (23±0.72°C) and ambient 2 (24±0.60°C) still stabilized below the critical threshold for cell viability. In the ‘Closed’ configuration, temperatures were consistently higher, yet they remained within acceptable limits, suggesting that passive heat dissipation mechanisms were sufficient to maintain a safe operational environment for the duration of the experiments.

### Vibration Characterization

Experimental characterization of the bioreactor’s vibrational dynamics, supplemented by COMSOL simulations, confirmed operational stability across the frequency spectrum of interest. The simulations indicated no resonant frequencies within the operational range, with the first mode of vibration occurring at 929 Hz (**Figure 3**), presenting minimal the risk of resonance interference. Empirical tests validated the simulation, demonstrating effective vibrational isolation at operational frequencies, with X/Y and Z/Y acceleration ratios remaining below 0.13 at lower frequencies. As frequency increased, a corresponding rise in these ratios was observed, suggesting higher transference of vibration to non-targeted axes. **Tables 1** and **2** encapsulates these findings, showing the bioreactor’s performance at various frequencies and accelerations, with a clear delineation between low and high-frequency behavior.

**Table 1:** Acceleration ratio in each non-targeted axes to the acceleration of target direction, highlighting the relative distribution of vibrational energy across all axes. . The bioreactor was subjected to vibration of 0.3g peak-to-peak across the entire 60 Hz to 500 Hz frequency range.

| Frequency (Hz) | X/Y | Z/Y |
| --- | --- | --- |
| 60 | 0.06 | 0.06 |
| 70 | 0.06 | 0.13 |
| 80 | 0.06 | 0.13 |
| 85 | 0.06 | 0.13 |
| 90 | 0.06 | 0.13 |
| 95 | 0.06 | 0.10 |
| 110 | 0.06 | 0.13 |
| 130 | 0.06 | 0.13 |
| 150 | 0.06 | 0.10 |
| 200 | 0.13 | 0.16 |
| 250 | 0.13 | 0.16 |
| 300 | 0.13 | 0.19 |
| 500 | 0.14 | 0.19 |

**Table 2:**
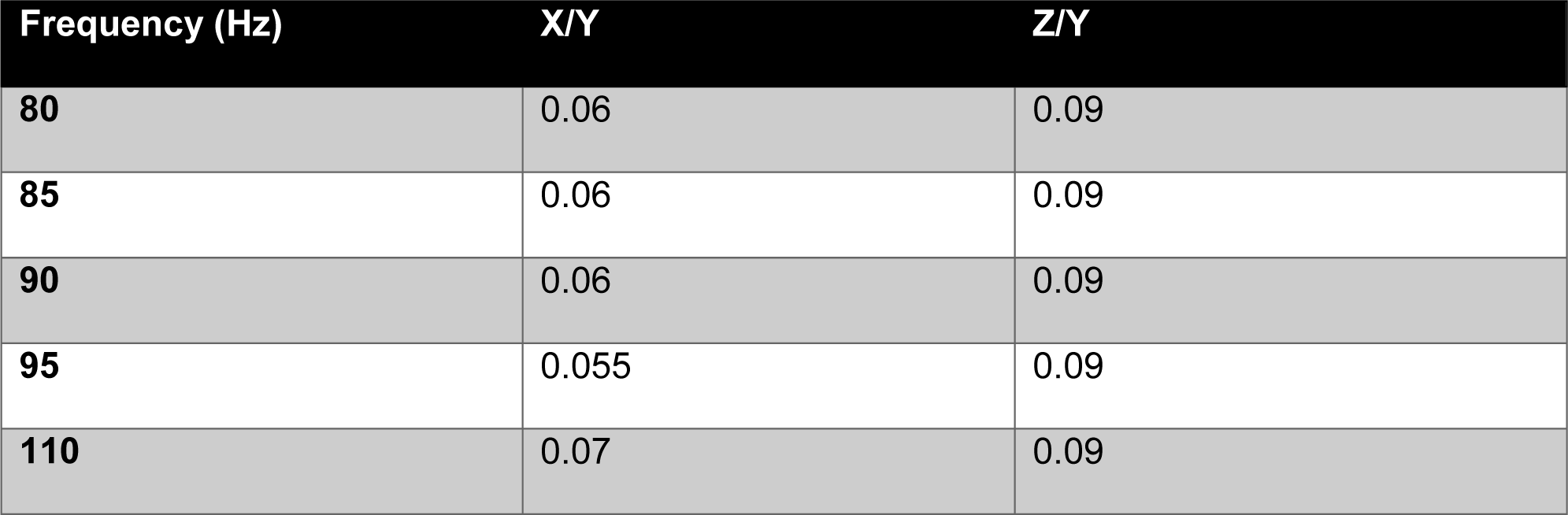
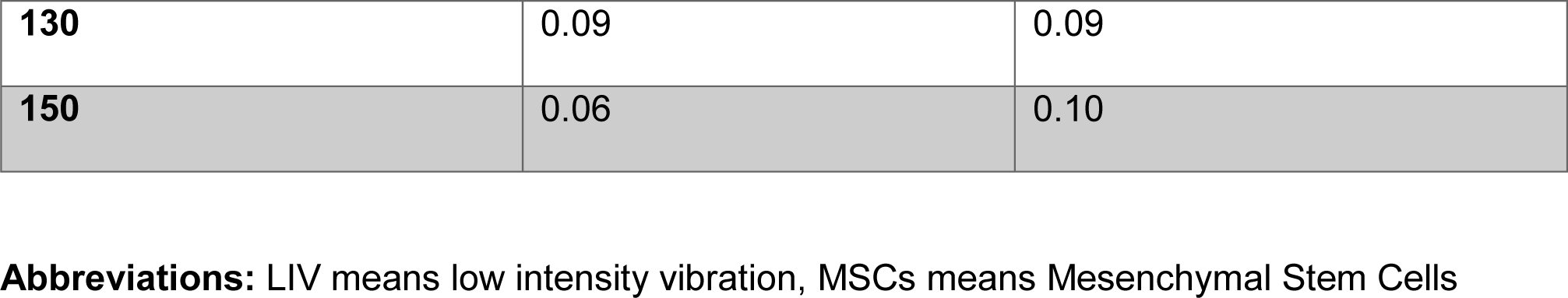
Acceleration ratio in each non-targeted axes to the acceleration of target direction, highlighting the relative distribution of vibrational energy across all axes. . The bioreactor was subjected to vibration of 0.7g peak-to-peak limited to frequencies between 80 Hz and 150 Hz due to system constraints.

### Scaffold viability

The XTT assay results indicated no significant difference in metabolic activity between the vibrated (+LIV) (p=0.9973) and non-vibrated (-LIV) (p>0.9999) samples from day 1 (D1) to day 4 (D4). Also, from D1 to D4, the metabolic activity of cells on the new scaffold material slightly increased in both vibrated (D1 mean: 1.4584 to D4 mean: 1.5453) and non-vibrated (D1 mean: 1.4584 to D4 mean: 1.4986) conditions. This showed that the BioMed Clear resin used in the scaffolds was biocompatible.

For a more direct observation of cell viability, Live/Dead imaging was employed. The imaging results, presented in **Figure 4c-f** and (**Supplementary Figure 4**), visually demonstrate the proportion of live (green) to dead (red) cells, providing a qualitative assessment of cell health. The pictures show that there are mostly living cells in both +LIV and -LIV conditions. There does not seem to be an increase in cell death after LIV was added or when the new material for the scaffold was added. The comparative analysis of the live/dead ratio further supports the conclusion that the new scaffold material, in conjunction with the LIV bioreactor, maintains a conducive environment for cell growth and viability.

### Diffusion Study

The diffusion characteristics within the CubeWells were initially modeled using COMSOL simulations to establish an efficient media exchange protocol. According to the simulations illustrated in **Figure 5b-c**, we observed that by employing a pumping rate of 1mL/min, approximately 80% of the volume of CubeWell would be occupied by fresh media during a span of 3 minutes. This rate of media replacement would result in total replacement of the media by the 5-minute mark. These findings informed the operational parameters for the media pump in subsequent in-vitro validation experiments.

In practice, the diffusion regime for the in-vitro experiment, visualized in **Figure 5a**, involved a 5-minute media pumping phase at the same flow rate, followed by a 15-minute recirculation period. The high-speed camera monitoring revealed a consistent media distribution pattern (**Figure 5d-e**), with a notable 55-second delay in reaching the center, bottom-left, and bottom-right locations compared to the simulation. Despite this delay, both the simulation and the experimental observations confirmed that 100% media exchange was accomplished within the 5-minute duration. The similarly between the predicted and actual diffusion patterns validates the simulation’s utility in setting practical experimental conditions and underscores the system’s capability to maintain cell viability by preventing prolonged exposure to stagnant media.

### Conclusions and Discussion

This study presented a wide-ranging characterization of a LIV bioreactor designed for microgravity conditions, such as those on the ISS. The research focused on validating the bioreactor’s mechanical and biological performance, ensuring its compatibility with the stringent requirements of space missions.

The selection of the linear guide was critical for the bioreactor’s performance. Our results indicated that the Schneeberger NKL 2-95 linear guide provided consistent one-axis acceleration, crucial for minimizing disturbances during vibration. Despite its larger size, it demonstrated superior performance in maintaining the desired vibrational profile with minimal cross-axis acceleration, as compared to the Igus N Prism guide and the Misumi linear guide. The Misumi linear guide, previously used in the laboratory, served as an effective benchmark, confirming the reliability of the Schneeberger guide under the test conditions.

Thermal characterization within the CubeLab environment demonstrated that the system could sustain a stable temperature profile, with variations within ±0.80°C over a 25-minute operational period. Notably, while the controller exhibited an increase in temperature, the duration of operation and the volume of air within the CubeLab were such that the rise in the overall ambient temperature did not create any problems in terms of either cell viability or piezoelectric controller’s operation range. The CubeLab’s exterior design proved effective in dissipating heat from the controller, thereby minimizing thermal transfer to the biological samples. This indicates that the bioreactor’s thermal management is adequate for the tested operational conditions without the need for active cooling systems. Nonetheless, the design accommodates the integration of minimal cooling solutions for experiments with cooler ambient requirements, ensuring adaptability for a broad spectrum of research needs.

The vibrational characterization of the bioreactor was approached with a strategy that combined predictive COMSOL simulations with empirical testing. The simulations provided theoretical assurance that the bioreactor’s design avoids resonant frequencies within the operational range, with the first resonant mode predicted at 929 Hz. While empirical testing did not directly confirm this resonant frequency, it did reveal an increase in the ratio of non-targeted to targeted axis accelerations at higher frequencies. This trend suggests that the bioreactor’s actual vibrational isolation performance aligns with the simulation outcomes, especially since no resonance was observed during the tested frequency range.

The increase in acceleration ratios at higher frequencies highlights the need for careful monitoring of vibrational behavior as operational frequencies approach the upper limit. Still, the bioreactor met the goals for vibration isolation, showing that it could keep accelerations under control on the main axis and prevent transference to the non-vibrated axes. This finding is crucial for microgravity research applications, where the fidelity of mechanical stimuli delivery to biological samples is of utmost importance.

The simulation’s finding that there is no resonance during routine operations, along with empirical evidence of vibration isolation, points to the bioreactor as a promising tool for spaceflight experiments. It is prepared to provide a stable environment for the study of microgravity’s effects on biological systems. Future design enhancements may focus on further reducing vibrational transfers at higher frequencies, leveraging the insights gained from this study to refine the bioreactor’s performance.

The evaluation of the scaffold’s biological performance within the LIV bioreactor demonstrated a slight increase in metabolic activity by day four, with a marginally higher increase observed in the vibrated (+LIV) samples. This increase, although subtle, hints at a potential positive interaction between mechanical stimulation and cellular activity over time with the use of the new scaffold material. The use of SLA-printed scaffolds across all samples ensures a consistent comparison base, highlighting the specific effects of LIV on cell viability.

The BioMed Clear resin scaffolds, employed in both vibrated and non-vibrated conditions, have shown promising compatibility for extended cell culture applications. This compatibility is crucial for future microgravity studies, where material behavior under prolonged exposure to LIV can significantly influence experimental outcomes. The data suggests that these scaffolds can support cell growth and maintain viability, which is essential for the success of long-duration biological experiments in space. Further research, with an extended timeline, will be instrumental in understanding the full scope of LIV’s effects on cellular systems in microgravity. The robustness of the SLA-printed scaffolds for intricate designs also opens avenues for more complex tissue engineering applications in space, where precise geometric control is crucial. The findings thus far provide a solid foundation for the extended use of these materials and methods in microgravity research.

Diffusion studies, supported by COMSOL simulations and in-vitro experiments, verified the CubeWell system’s media exchange efficiency. The congruence of simulation and experimental outcomes validated the media pump’s operational parameters, ensuring timely and uniform cell nourishment without prolonged exposure.

In conclusion, the contributions of this research are versatile, addressing previous limitations in space research and providing a comprehensive solution that enhances the scope of biological experimentation in microgravity. The insights gained from this work are expected to have a profound impact on the health and safety of astronauts by informing the design of future space missions and the development of countermeasures against the adverse effects of living in space.

## Acknowledgements

This study was supported by, AG059923, P20GM109095, P20GM103408, NSF1929188 and, NSF 2025505

## Data Availability

The datasets used and/or analyzed during the current study are available from the corresponding author on reasonable request.

## Contributions

Omor Khan: concept/design, data analysis/interpretation, manuscript writing

Will Gasperini: concept/design, data analysis/interpretation

Chess Necessary: concept/design

Zach Jacobs: concept/design

Sam Perry: concept/design

Paul Kuehl: concept/design

Maximilien DeLeon: concept/design, data analysis/interpretation, manuscript writing

Kendall Nelson: concept/design, data analysis/interpretation

Paul Gamble: concept/design, final approval of manuscript

Anamaria Zavala: data analysis, concept/design, final approval of manuscript

Sean Howard: data analysis, concept/design, final approval of manuscript

Mary Farach-Carson: data analysis, concept/design

Elizabeth Blaber: data analysis, concept/design

Danielle Wu: data analysis, concept/design

Aykut Satici: concept/design, data analysis/interpretation, financial support, manuscript writing, final approval of manuscript

Gunes Uzer: concept/design, data analysis/interpretation, financial support, manuscript writing, final approval of manuscript

## Supplementary Figures

**Supplementary Figure 1:**
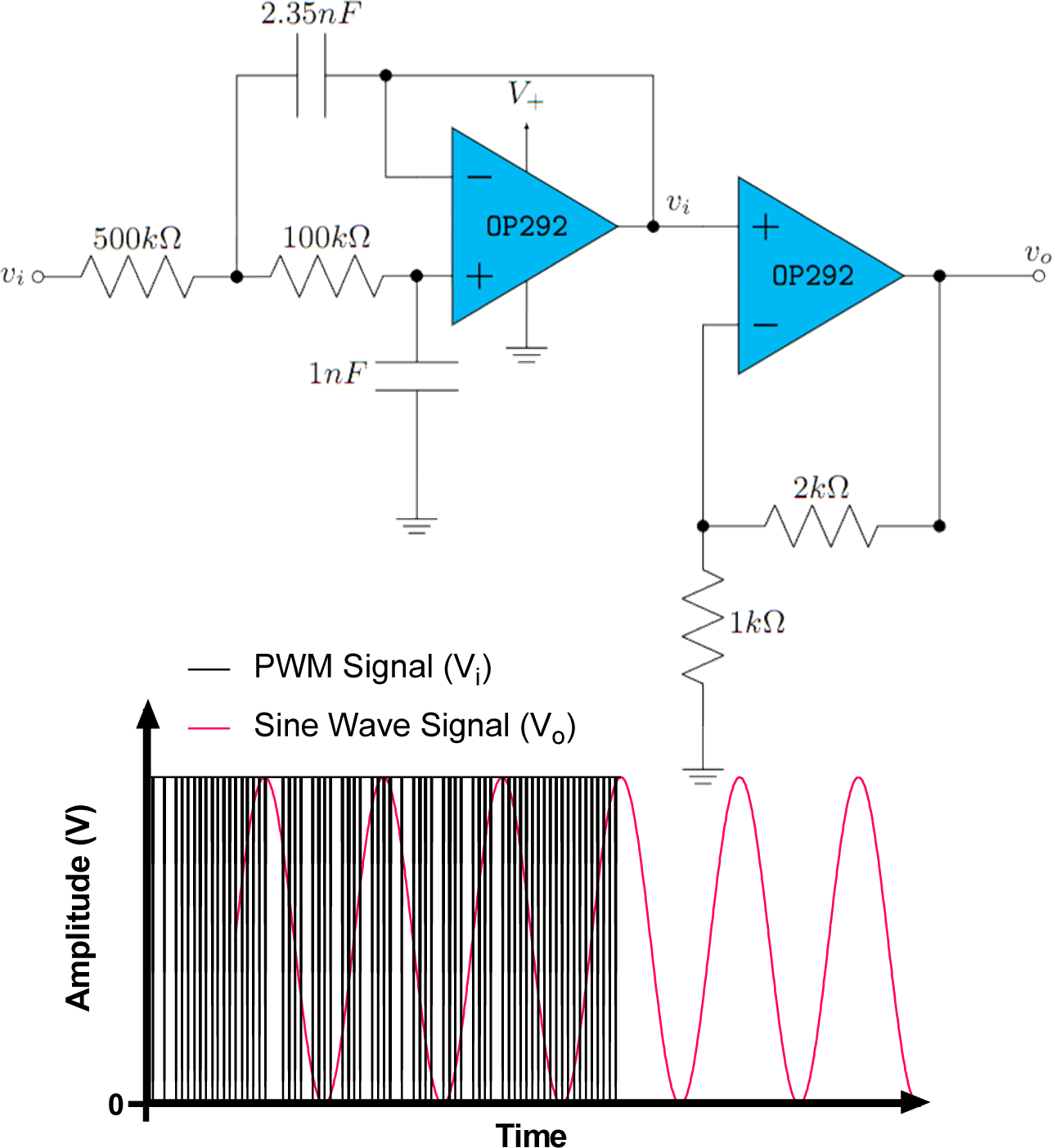
Custom circuit diagram (top) to generate sine wave signal from Teensy 4.0 microcontroller. PWM signal (square wave) coming from the microcontroller is filtered by the demodulating circuit to get a sine wave output that will be used as an input signal for the LIV.

**Supplementary Figure 2:**
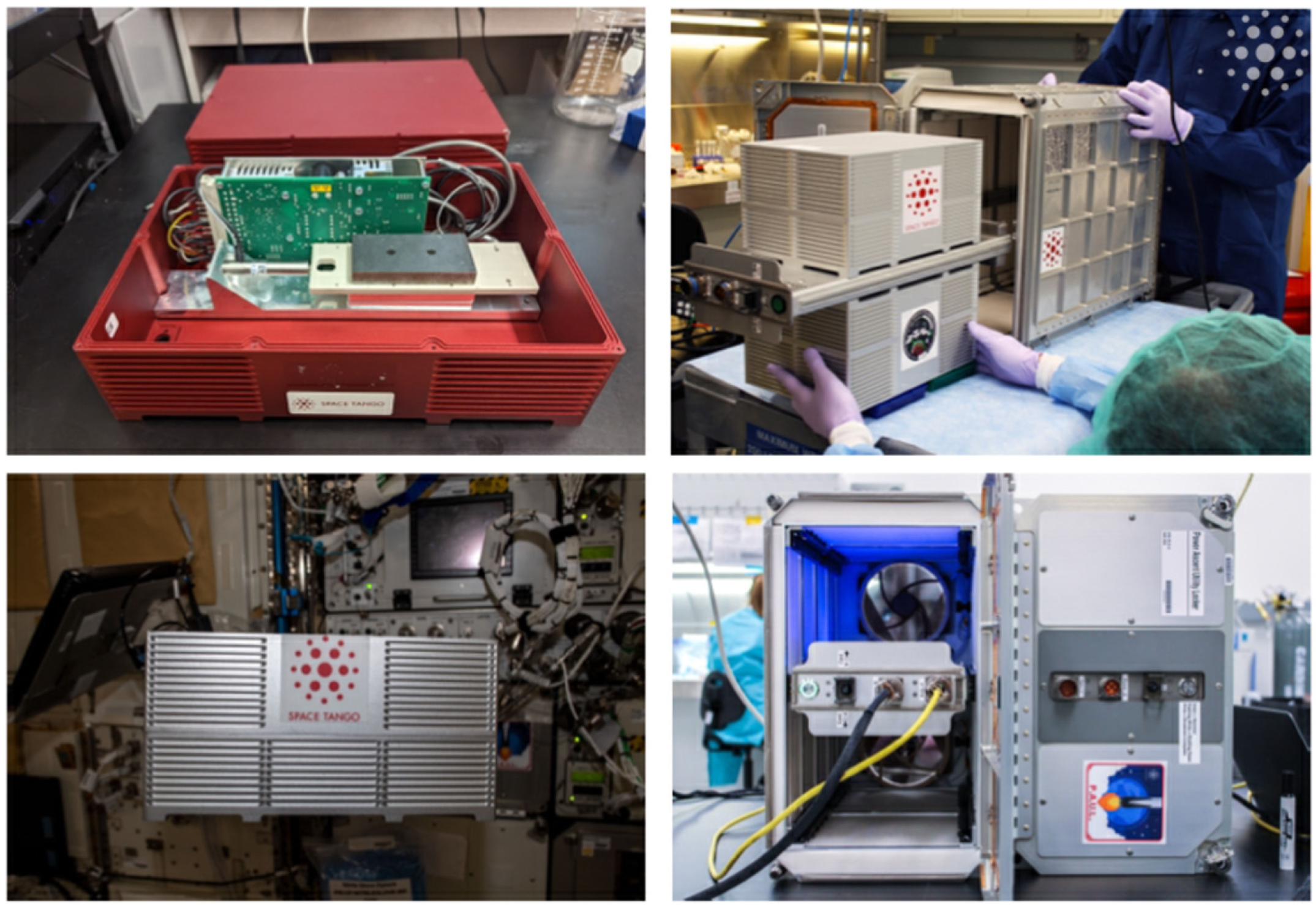
CubeLab (top and bottom-left) facility that will house the vibrational bioreactor and the biological samples. The PAUL (top and bottom-right) facility designed to house and provide mechanical, electrical, and network interface from the outside.

**Supplementary Figure 3:**
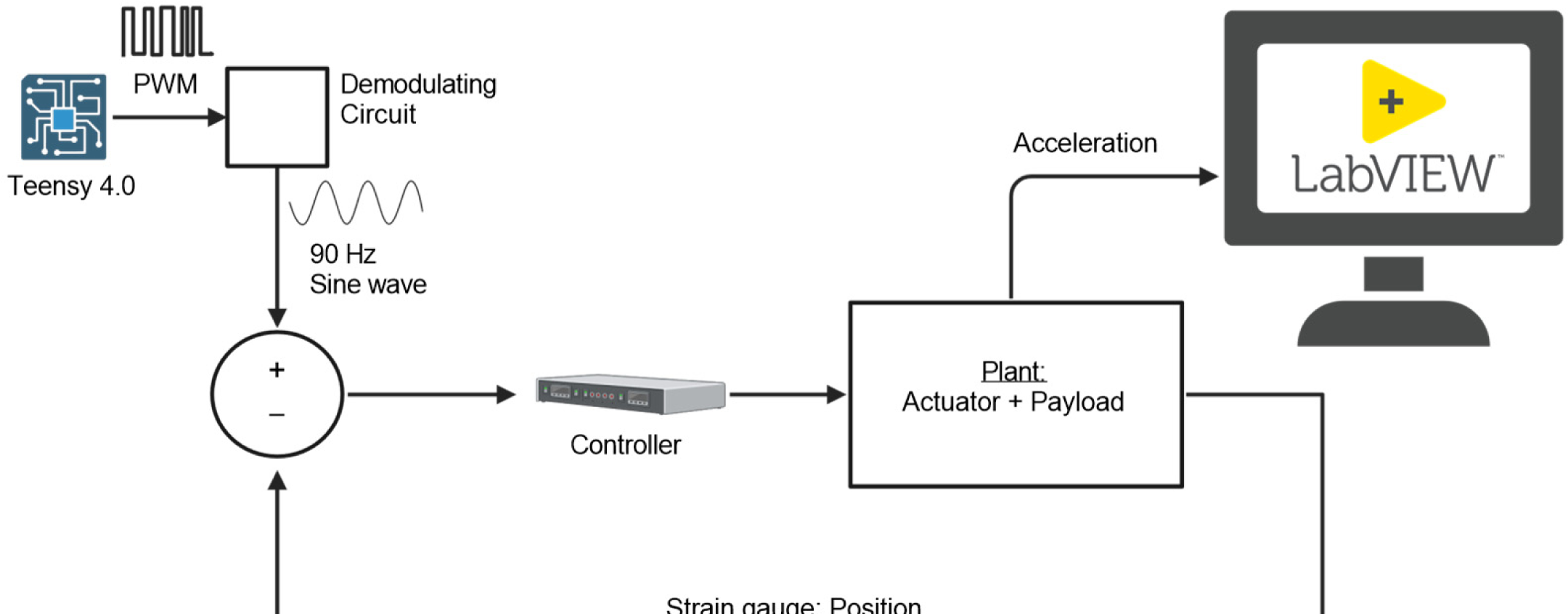
Experiment schematic of the LIV regimen from signal generation to data acquisition and visualization in LabVIEW.

**Supplementary Figure 4:**
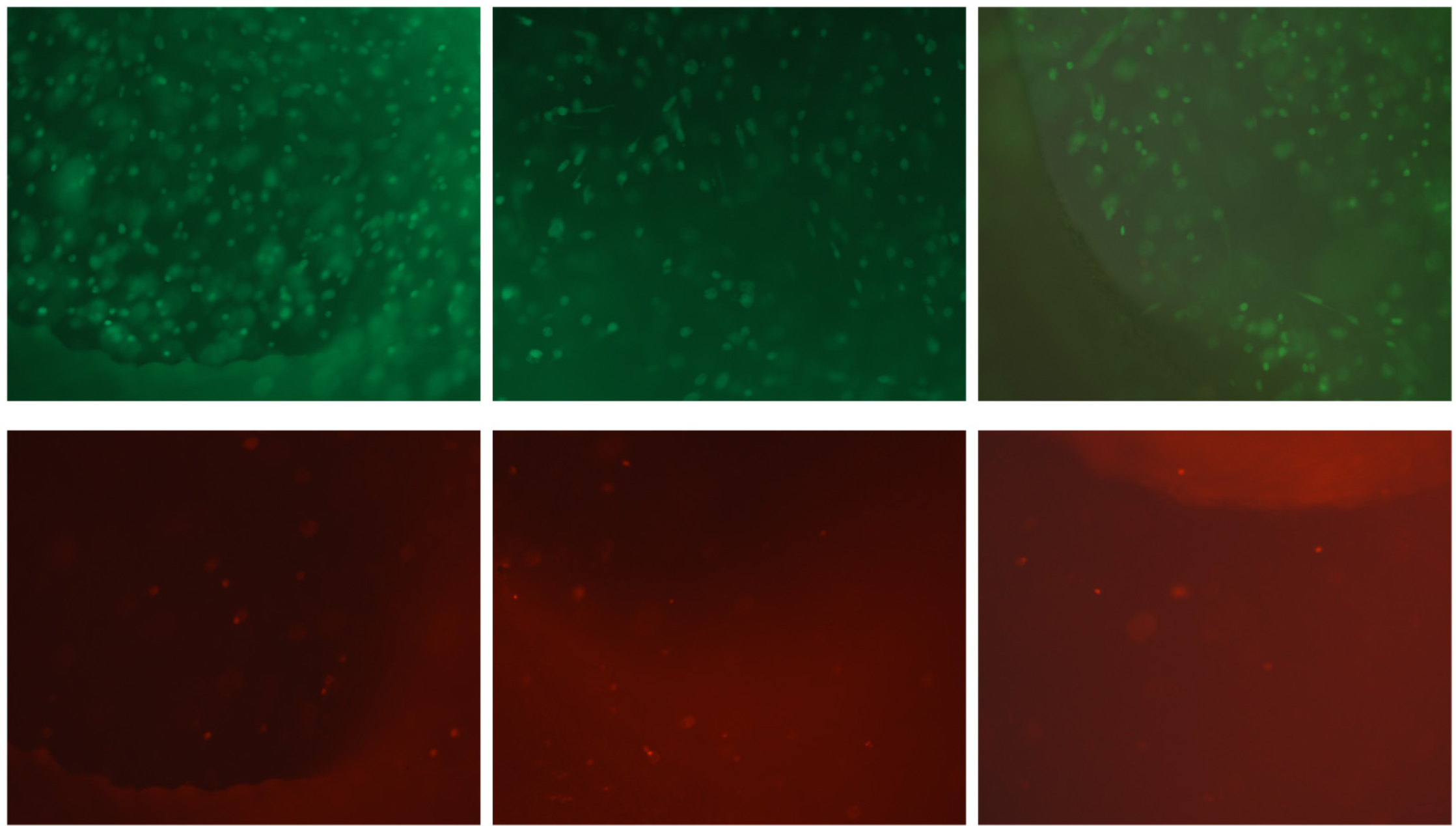
Additional Live/Dead assay images across various scaffold locations and samples, demonstrating a predominance of live cells over dead ones. These images further validate the biocompatibility and cell-supportive nature of the scaffold materials used in the study.

**Supplementary Figure 5:**
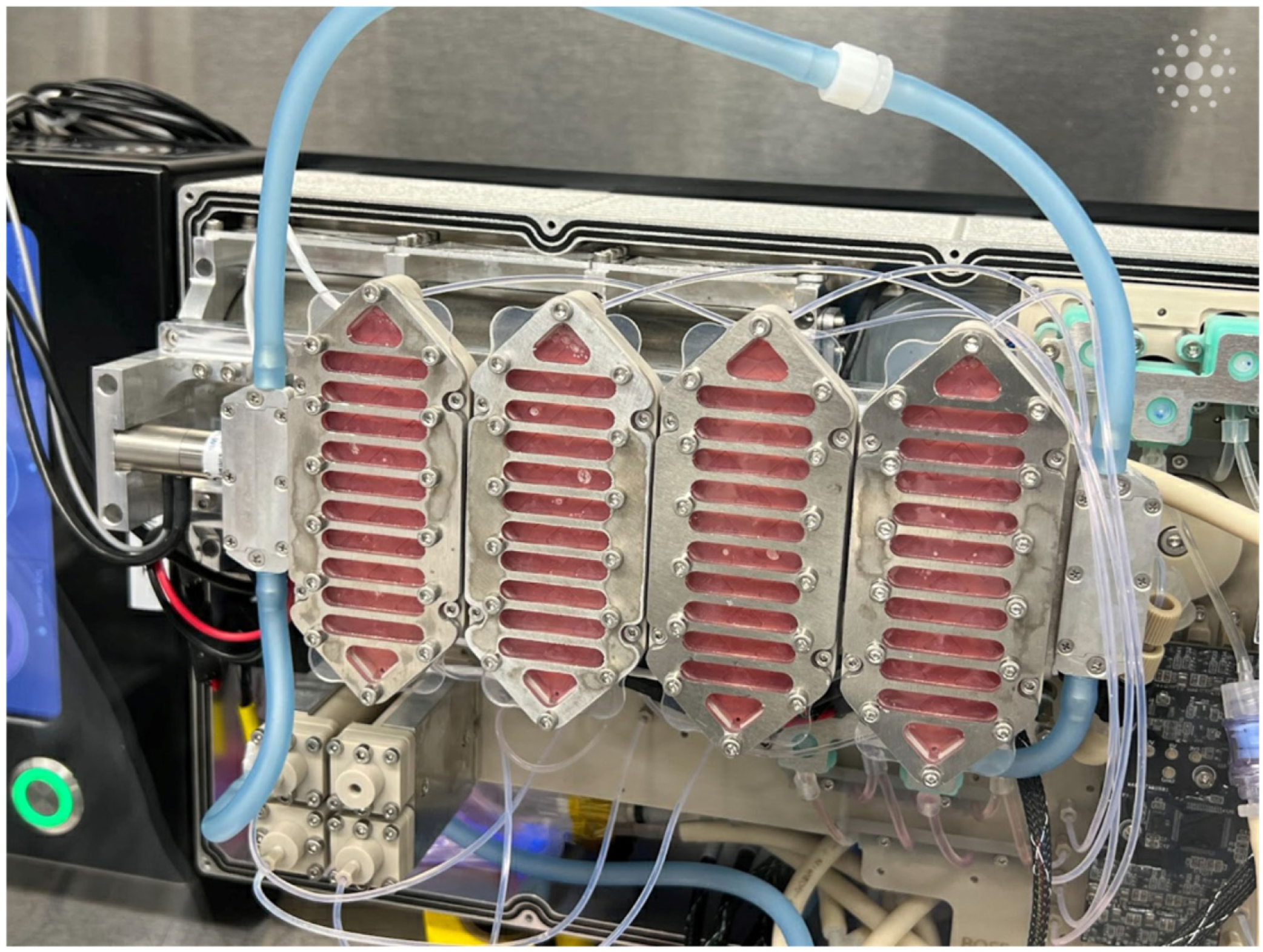
CubeWells within the CubeLab module, showcasing the custom-designed well plates developed for holding hydrogel-encapsulated scaffolds. The design incorporates a secure lid system to prevent sample spillage during spaceflight, with scaffolds protected by a biocompatible PDMS layer, further sealed by a metal lid for added safety and integrity during experiments in space.

